# Mismatch of the morphology model is mostly unproblematic in total-evidence dating: insights from an extensive simulation study

**DOI:** 10.1101/679084

**Authors:** Seraina Klopfstein, Remo Ryer, Mario Coiro, Tamara Spasojevic

## Abstract

Calibrating the molecular clock is the most contentious step in every dating analysis, but the emerging total-evidence dating approach promises increased objectivity. It combines molecular and morphological data of extant and fossil taxa in a Bayesian framework. Information about absolute node ages stems from the inferred fossil placements and associated branch lengths, under the assumption of a morphological clock. We here use computer simulations to assess the impact of mismatch of the morphology model, such as misspecification of character states and transition rates, non-stationarity of the evolutionary process, and extensive variation of evolutionary rates among branches. Comparisons with published datasets suggest that, at least for evolutionary rates typically observed in discrete morphological characters, the total-evidence dating framework is surprisingly robust to these factors. We show that even with relatively low numbers of morphological characters sampled, extensive model mismatch is mostly irrelevant for the performance of the method. The only exception we found are cases of highly asymmetric state frequencies and thus transition rates, but these can be accounted for by appropriate morphology models. In contrast, we find that the temporal scope of fossil sampling has a major impact on divergence time estimates, with the time signal quickly eroding if only rather young fossils are included in an analysis. Our results suggest that total-evidence dating might work even without a good understanding of morphological evolution and that study design should instead focus on an adequate sampling of all relevant fossils, even those with highly incomplete preservation.

## Introduction

The estimation of absolute timescales based on molecular phylogenies is one of the most contentious issue in contemporary systematics, with age estimates for particular groups varying by up to an order of magnitude depending on the applied methodology (Donoghue and Benton 2007; Inoue, et al. 2010). The traditional “node dating” approach has been exposed as subjective through sensitivity analyses, which showed that drastically different age estimates can result from the analysis of the same fossil sample (Misof, et al. 2014; Tong, et al. 2014; Warnock, et al. 2012). The main issue with node dating is that while the oldest fossil of a group can provide a minimum age for its ancestral node, this is not sufficient for informing divergence time estimates; instead, more informative node age priors are implemented that use distributions with mean and/or maximum values, but these are often hard to justify (Fig. 1). Moreover, the interaction between different node priors and the tree prior often leads to effective calibration densities that diverge substantially from the ones specified by the user (Brown and Smith 2018), paradoxically reducing the advantage of implementing more fossil information. Node calibrations have furthermore been exposed as highly problematic if they happen to be close to strong shifts in evolutionary rates, which has been suggested as an important reason for the conflict between molecular date estimates and interpretations of the fossil record in major groups (angiosperms: Beaulieu, et al. 2015; Brown and Smith 2018; Coiro, et al. 2019; mammals: Phillips and Fruciano 2018).

**Figure 1.**
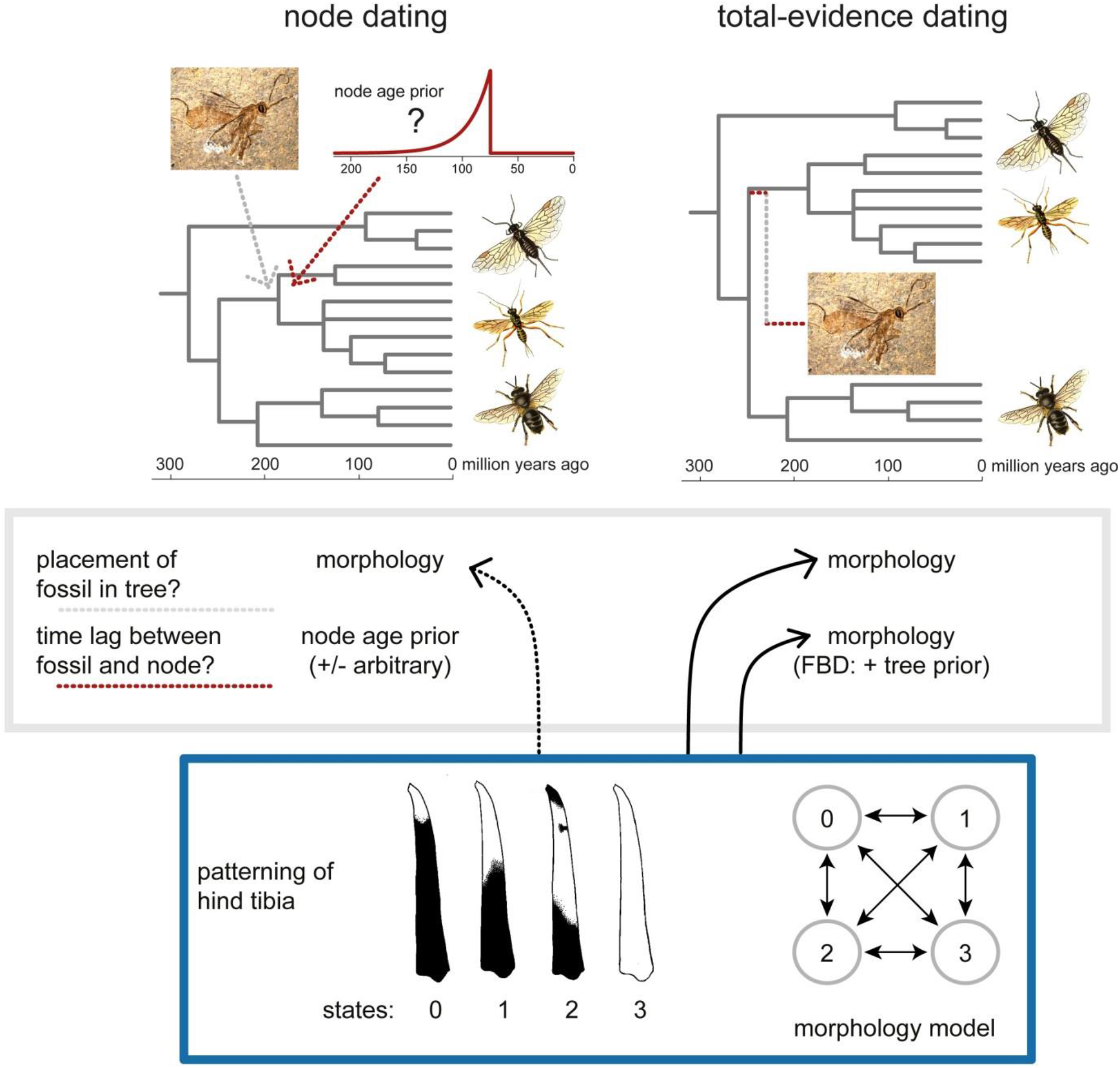
Different sources of information used for fossil placement and to estimate the time lag between fossil and associated node. In node dating, morphological data is used either in a qualitative (i.e., by invoking synapomorphic characters) or quantitative way (i.e., in a morphological phylogenetic analysis) to place a fossil in the tree of extant taxa. The node-age prior, which is set on the node below the likely placement of the fossil, is chosen in a largely arbitrary fashion. In total-evidence dating, fossil placement is directed by morphology in connection with a morphological model, and uncertainty in fossil placement can be integrated over. The time lag between the fossil and its subtending node is based on the morphological data. In the case of an informative tree prior such as the fossilized-birth-death prior, information about branch lengths also comes from the tree prior.

An alternative approach, called “total-evidence dating” (TED; Fig. 1), includes fossils as terminals directly in the analysis. This allows to integrate over the uncertainty in the phylogenetic placement of these fossils, which can be extensive (e.g., Klopfstein and Spasojevic 2019; Ronquist, et al. 2012a). The information for fossil placement in this approach comes from a matrix that is compiled by coding morphological characters both for fossil and extant taxa. This morphological matrix is then analysed alongside molecular data for the extant taxa, applying models for the morphology partition that are similar to those used for DNA characters (Lewis 2001). Divergence time estimates then depend both on the placement of the fossils in the tree and on the lengths of the branches leading to them; these are in turn informed both by the morphological data and by the chosen tree prior. Two variants are currently in use as tree priors in TED analyses: the uniform tree prior on both topology and branch lengths (Ronquist, et al. 2012a), which acts as an uninformative prior, and a parametric tree prior that is based on the fossilized birth-death (FBD) process (Heath, et al. 2014).

Recent studies have shown that the FBD tree prior can be highly informative and potentially overrule any information about branch lengths that might have been present in the morphological partition (Brown and Smith 2018; Parins-Fukuchi and Brown preprint; Ronquist, et al. 2016). This has a clear implication for the experimental design of TED studies: if the fossil record of a group is rather extensive and regular enough for it to be modelled adequately, but interpretable morphological characters are few, then it would seem reasonable to rather put our trust in an inferred fossilization process than in morphological branch lengths; the opposite might be true for groups with a sparse and irregular fossil record, but reasonable morphological preservation. In any case, assumptions in TED analyses that influence the choice of the tree prior should be made explicit and marginal priors published alongside posterior age estimates (Brown and Smith 2018). Currently, we lack a good understanding of how model mismatch impacts both approaches – mismatch of the fossilization model for TED analyses that rely largely on FBD-informed branch lengths versus mismatch of the morphology model for those using the uniform tree prior. The former was the subject of two previous simulation studies (Luo, et al. 2019; O’Reilly and Donoghue 2019), while we here focus on the latter aspect.

While some studies that performed TED reported a good fit between the results of the analysis and the expectations based on the fossil record (Dornburg, et al. 2015; Gavryushkina, et al. 2017; Near, et al. 2014; Ronquist, et al. 2012a; Wood, et al. 2013), others suspected that their divergence time estimates might be too old (Arcila, et al. 2015; Beck and Lee 2014; Lee, et al. 2014). Besides a general lack of reporting of marginal priors on node ages in most of these studies (Brown and Smith 2018; Parins-Fukuchi and Brown preprint), almost all of them combined TED with the FBD tree prior and/or at least one informative node-age prior, which makes it difficult to narrow down the reasons for these suspected biases. What has been suspected repeatedly as the likely culprit is a mismatch between the data and the morphology model, especially due to an inability to define character states and transitions between them with sufficient biological reality (Beck and Lee 2014). Observed character states in morphology are believed to be difficult to delimit in an objective way, and transitions between them might not adequately reflect the underlying biochemical and developmental processes (Tarasov 2019). Furthermore, morphological characters are generally assumed to have a low clocklikeness (Beck and Lee 2014; Lee 2016; Lee, et al. 2014). The pace of morphological evolution is usually thought to be influenced by episodic speed-ups, associated for instance with a change in the life-history of a lineage (which would impact the evolutionary rate of the entire genome, e.g., Thomas, et al. 2010), or with divergent selection on a set of morphological characters after ecological shifts (leading to heterotachy; Goloboff, et al. in press). This has led some authors to reject the idea of a common mechanism acting on all morphological traits of an organism (Goloboff, et al. in press). The lack of a theoretical grounding of the morphological clock hypothesis has also been cited (Parins-Fukuchi and Brown preprint), while the molecular clock assumption is grounded in the neutral hypothesis of molecular evolution (Kimura 1968). However, nucleotide and amino acid sites that follow neutral or nearly-neutral evolution typically evolve rather fast because they are not under purifying selection, while molecular dating studies that address deep timescales rely heavily on slowly-evolving sections of DNA for which it is difficult to claim neutral evolution. The conceptual difference between the molecular and the morphological clock might thus not be that pronounced after all, and quantitative comparisons between the clocklikeness of morphological versus molecular partitions are scarce (but see, e.g., Beck and Lee 2014; Ronquist, et al. 2012a). Furthermore, a strict-clock model is likely inadequate also for most molecular partitions (Drummond, et al. 2006), and the question is thus not whether a morphological clock exists at all, but rather how clocklike morphology needs to evolve in order for TED to be able to extract sufficient time information from it.

The standard model for morphological data was elaborated by Lewis (2001), who proposed using a Markov model with equal state frequencies and equal rates (“Mk model”), with the number of states (“k”) chosen according to the observed number of states in each character. This symmetric model is identical to the Jukes-Cantor model for nucleotides in the case of k = 4, and it circumvents the issue of arbitrary state labels in morphological data. As with models for nucleotide and amino-acid evolution, it assumes stationarity, reversibility and homogeneity of the process on the phylogeny in question (Jermiin, et al. 2008), as well as independent evolution of each character. Lewis (2001) already proposed important extensions of this simple model, such as among-character rate variation (modeled for instance under a discretized gamma-distribution as for molecular data; Yang 1994), which has proven to be a vital aspect in many morphological datasets (Harrison and Larsson 2015; Wagner 2011). Second, Lewis (2001) suggested a way to account for asymmetry in state frequencies and thus in transition rates, an improvement that only recently started to attract attention (Klopfstein and Spasojevic 2019; Wright, et al. 2016). Furthermore, it is straightforward to incorporate non-stationarity and thus directional evolutionary patterns by introducing separate state frequencies at the root (Klopfstein, et al. 2015). Other types of model mismatch might be more difficult to overcome, such as pronounced heterotachy (Goloboff, et al. in press) or non-homogeneity of the evolutionary process (Jayaswal, et al. 2011; Jermiin, et al. 2008). Alekseyenko et al. (2008) proposed a combination of an Mk-like model and a binary stochastic Dollo-model based on the Poisson process, thus allowing for two types of transitions within a single character; this might be an adequate depiction of multi-state characters that include an “absent” state and several modifications of the “present” state, as in the famous example of a tail with different colours. Tarasov (2019) went even further by proposing hidden-state Markov models that reflect the mismatch between our character concepts and the underlying evolutionary process. All these models promise to lead to more robust phylogenetic inference from morphological data and improve our understanding of morphological evolution; however, it remains unclear how much model realism is in fact needed to obtain accurate divergence-time estimates in the TED framework.

We here perform an extensive simulation study to explore the limits of TED with respect to mismatch of the morphology model under the uniform tree prior. We simulate molecular and morphological datasets on two sets of trees of differing sizes (12 extant plus 4 fossil taxa, and 48 extant plus 16 fossil taxa). In these simulations we introduced all kinds of potentially detrimental biases, including high levels of homoplasy, misspecification of character states and transitions, and deviations from clocklike evolution in the morphology partition. We also explore the impact of different fossil sampling strategies on TED performance. Finally, we compare insights from our simulations with the characteristics of empirical morphological datasets used in TED studies, which allows us to make recommendations for future dating studies.

## Results

### Saturation of the morphology partition

Overall, total-evidence dating performed very well on our simulated datasets, retrieving the correct age of 100 million years in most of the replicates even for the smaller simulation trees with 12 extant and four fossil taxa (Fig. 2a). To examine the impact of saturation, we simulated morphological characters evolving at different evolutionary rates, which gave very similar results as a recent simulation study on phylogenetic informativeness (Klopfstein, et al. 2017): rates between 0.05 and 0.75 substitutions between root and tip performed well, as long as the characters were variable (Fig. 2b). If invariant sites were also sampled (as is the rule for molecular characters, but which is very difficult to do for morphology), then very low rates led to a loss of information. The inclusion or exclusion of autapomorphies in the matrices (without accounting for it in the analyses) did not have any discernible effect. Increasing the tree size strongly improved the robustness of total-evidence dating to high morphological rates, despite the increase in the actual number of changes on the full tree (and the related decrease in consistency and retention indices; results not shown). In fact, even at a rate of two changes between root and tips, all replicates on the larger tree recovered the true root age with very high precision (Fig. 2b).

**Figure 2.**
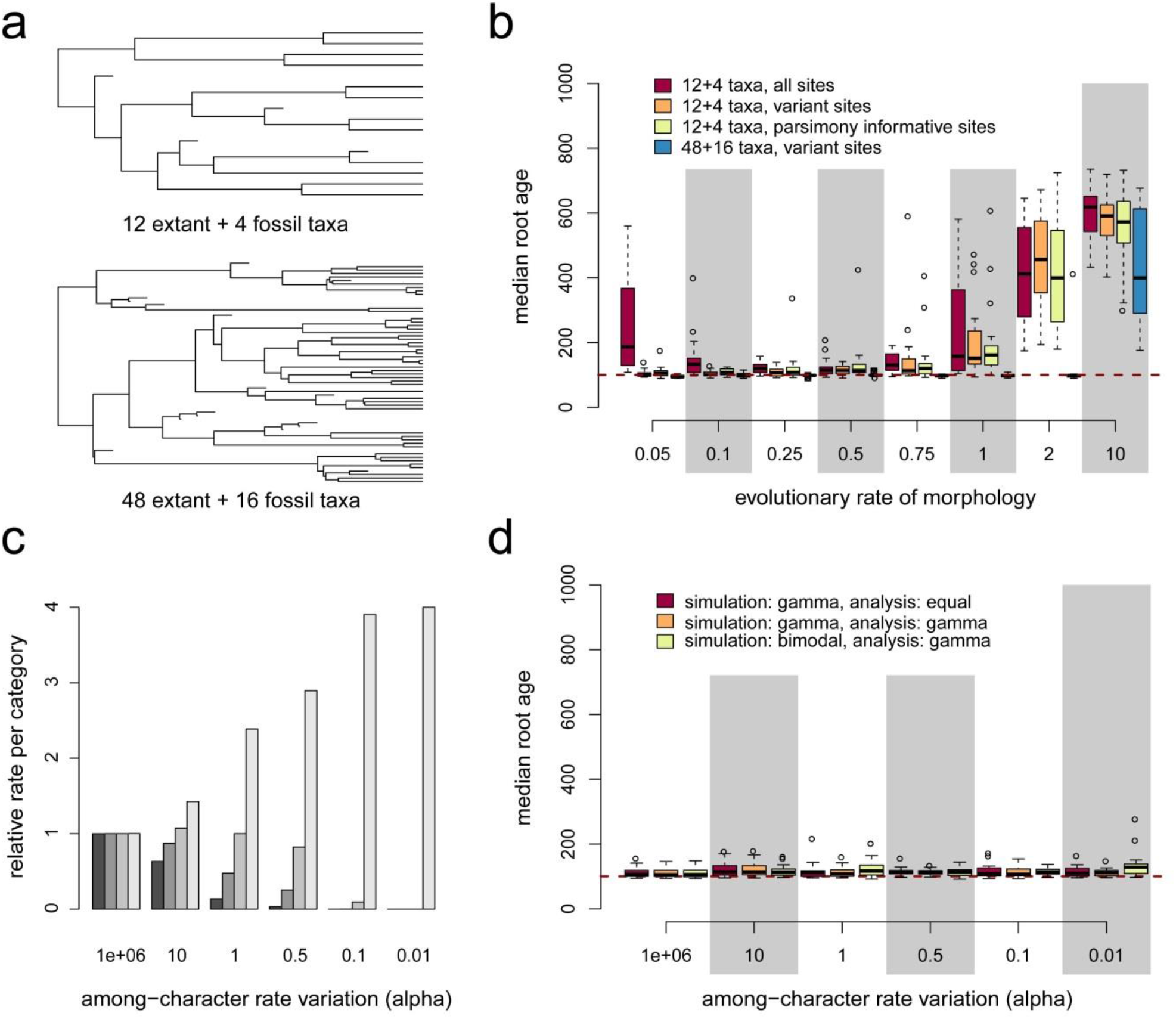
Simulation trees and analyses of evolutionary rate and among-character rate variation (ACRV). a) Two examples of trees simulated under the standard approach, with ¼ of taxa being fossils. Simulation trees have a fixed depth of 100 Ma and fossils are sampled evenly between 85 Ma and 15 Ma, with terminal branches leading to fossils being 5 Myr long. b) While the molecular partition was always evolving at a rate of 0.25 expected substitutions between root and tips (i.e., a rate of 0.0025 per Myr), the rate of the morphological partition is varied from 0.05 and 10. Boxplot depict the median root ages recovered over the independent replicates (20 for trees with 12 + 4 taxa, 10 for those with 48 + 16 taxa), and the dashed horizontal line indicates the true value (the clockrate prior implies a posterior of the root age that is very flat with a mean around 575 Ma). c) Relative evolutionary rate for each of the four simulated categories with increasing among-character rate variation. d) Impact of ACRV and misspecification of its model on root age estimates. Irregularities of the histograms in cases of low performance are due to the small number of replicates and should not be overinterpreted.

Rate variation among the morphological characters (ACRV) and lack of adequate modelling thereof during the analysis (Figs 2c and d) did not have a strong impact on total-evidence dating performance, even under very strong variation in the rates. This was irrespective of whether a model with equal rates was used during the analysis, or a matching or mismatching model that did allow for ACRV.

### Misfit of the Markov model

Mismatch of states and transition rates in the Markov model used for simulations versus analysis only showed a negative impact at very high evolutionary rates (Fig. 3). When characters were simulated under an ordered model (Fig. 3a), performance decreased with increasing evolutionary rate and with a larger number of ordered states, reflecting more pronounced model mismatch (Fig. 3b). Comparison of the two-states analysis (for which the ordered and unordered model are identical, i.e., there is no model mismatch) with the analyses of characters with three to ten states demonstrates that the erosion of the temporal signal mostly depends on the evolutionary rate, and not on model mismatch; only under extensive model mismatch (ten ordered states in the simulations) were rates of 1.0 or 2.0 expected changes per 100 Myr sufficient to perturb the analyses, while rates of 0.5 or lower always perform reasonably well (Fig. 3b).

**Figure 3.**
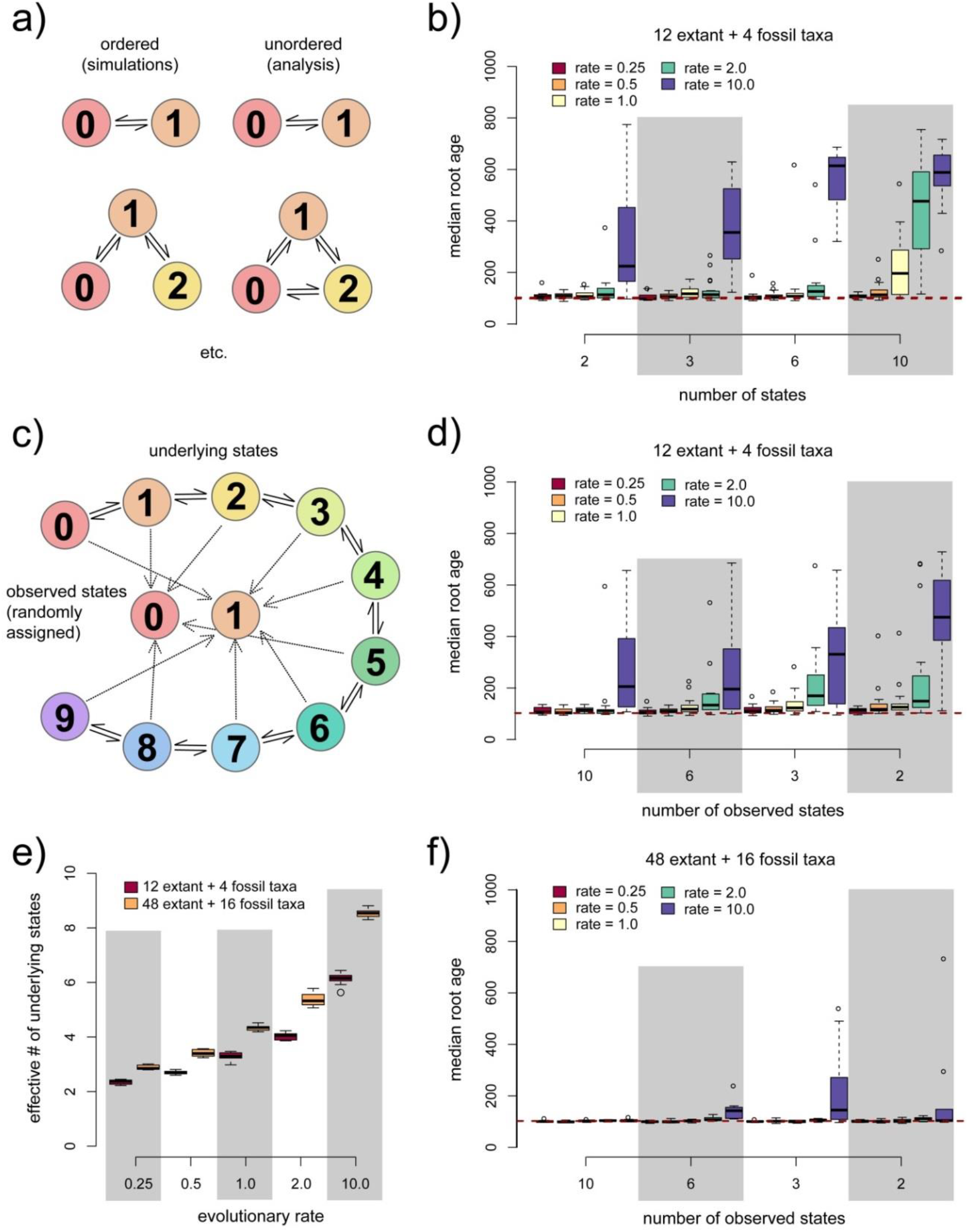
Misfit of character states and transitions between them in the Markov model. a) Depiction of the ordered versus unordered character transition model for two- and three-state characters. For two states, both models are equivalent; with increasing numbers of states, more of the possible transitions between states are not allowed under the ordered model, creating increasing extents of model mismatch. b) Results from analyses of ordered character data assuming the unordered model. Beware that the simulations of two-state characters show no model mismatch, thus reflecting the effect of the signal erosion due to the evolutionary rates only (compare to Fig. 2b). c) Depiction of the simulation procedure to mimic mismatches in both states and transition rates: characters were simulated under an ordered ten-state model and then randomly transformed to fewer and fewer observable states. d) Results from the transformed characters for trees with 12 extant + 4 fossil taxa. e) Effective number of underlying states resulting at the tips of the smaller versus the larger trees under different evolutionary rates. f) as d), but using trees with 48 extant + 16 fossil taxa. Irregularities of the histograms in cases of low performance are due to the small number of replicates and should not be overinterpreted.

When both character states and transitions between them were misspecified in the analyses versus simulations (Fig. 3c), the picture was very similar: performance of total-evidence dating decreased significantly only at rates above about 2.0 expected changes from root to tips (Fig. 3d), and this was even more the case for larger trees (Fig. 3f). A potential reason for this surprisingly high robustness of TED is that under low evolutionary rates, even if assuming that ten underlying states are transformed into just two observable states, the number of states that actually evolved on the tree is much lower than ten, leading to a lower effective loss of information than might be expected (Fig. 3e).

Asymmetric state frequencies pose a larger problem for TED (Fig 4). The simulations with equal state frequencies (50% each) showed the familiar picture, with rates below 1.0 performing very well already in smaller trees (12 extant taxa, Fig. 4b) and even more so in larger trees (48 extant taxa, Fig. 4c). Under moderate levels of asymmetry in character states (with character frequencies of 25% versus 75%), slow characters still performed rather well, but high asymmetry (90% versus 10%) led to a diminishing of the temporal signal even in very slowly-evolving characters (Fig. 4b). For larger trees, performance was a bit better for the lowest rates simulated (0.25 changes between root and tips), but rates higher than that already led to a strongly diminished performance (Figs 4b and c).

**Figure 4.**
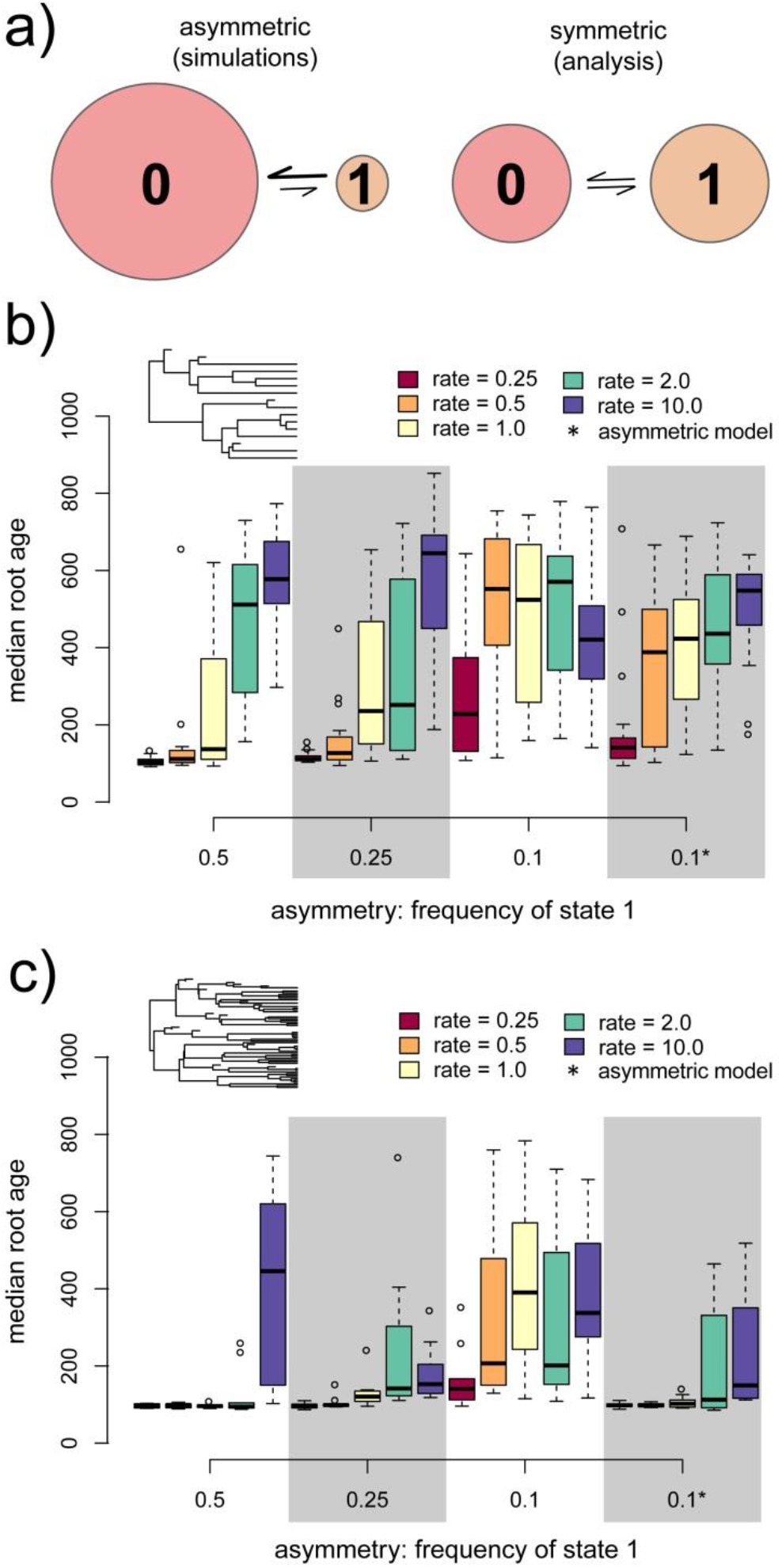
Asymmetric character states. a) Depiction of asymmetrical character states and transition rates in a two-state character. b) Results from the analyses of trees with 12 extant + 4 fossil taxa and symmetric characters (50% frequency in both states) versus characters simulated under different levels of asymmetry (25% versus 75% and 10% versus 90% state frequency). The analysis model assumed symmetric characters, except in the last column (marked as “0.1*”), where asymmetry was modelled under a symmetric Dirichlet distribution with a hyperprior (exponential with rate 10). Increasing levels of asymmetry lead to a decreasing performance of slowly-evolving characters, while performance is low throughout for fast characters. c) As b), but with trees with 48 extant + 16 fossil taxa; accounting for asymmetry in the analysis model here revoked the negative impact of asymmetry, except at high rates. Irregularities of the histograms in cases of low performance are due to the small number of replicates and should not be overinterpreted.

In contrast to asymmetry, non-stationarity in morphological evolution did not show a negative impact (Figs 5a and b). Even when all characters were assumed to evolve in a directional manner, i.e., when all 100 simulated characters started at the root of the tree in state ‘0’ and then evolved towards equal frequencies of states ‘0’ and ‘1’ (Fig. 5a), there was no difference in the performance of TED (although once more, it is obvious that characters evolving at rates higher than 1.0 expected changes between root and tips perform much worse than slow characters). This is the case even though the state frequencies differed considerably between the root, the fossils and the extant taxa, even at low rates (Fig. 5b).

**Figure 5.**
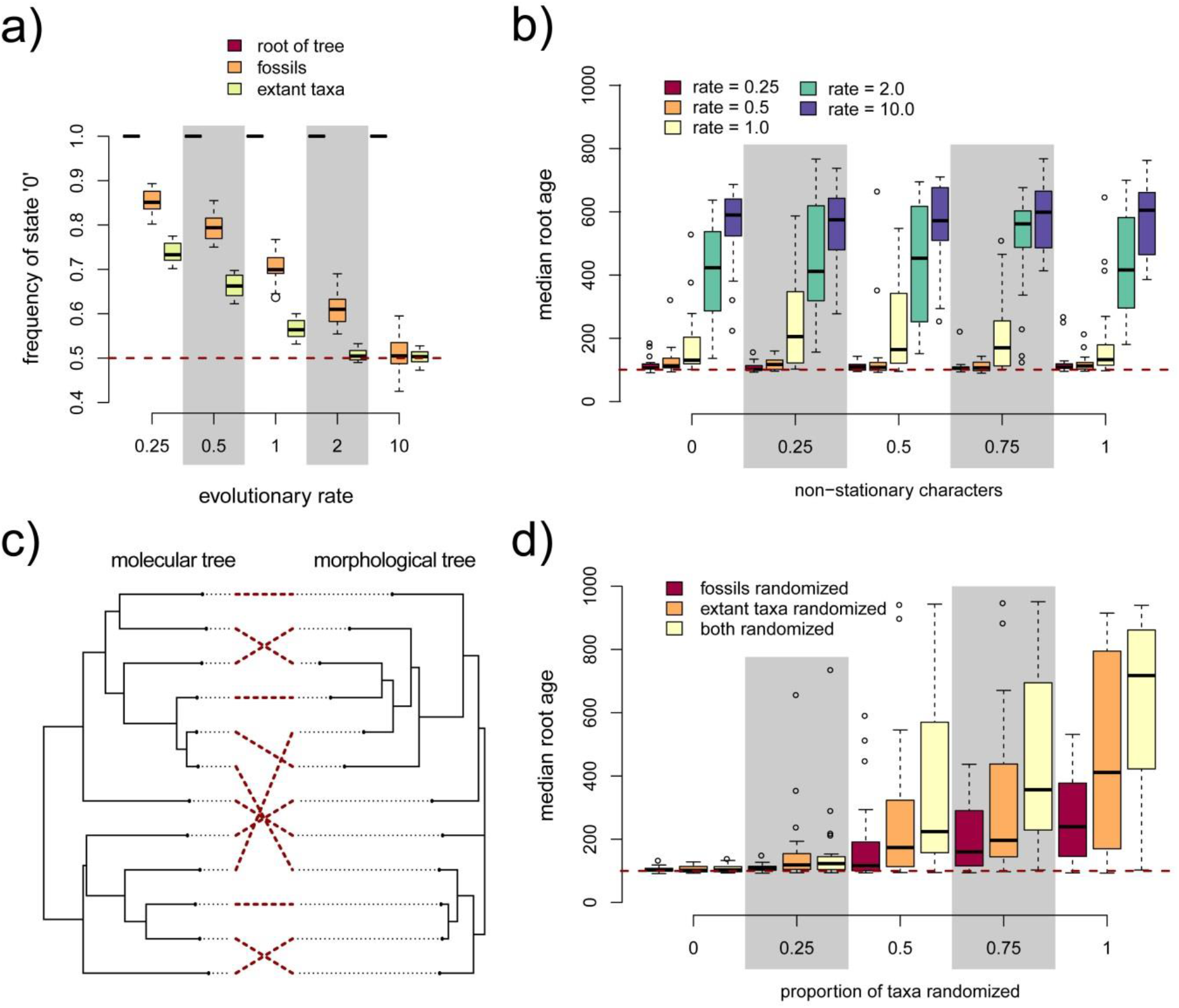
Non-stationarity and non-phylogenetic signals. a) Frequency of state ‘0’ as it resulted from the simulations of non-stationary characters. In all cases, the root frequency was assumed as 100%, while the stationary frequencies of both states are 50%. The average frequency across the fossils (aged between 85 and 15 Ma) differs in all but the fastest-evolving datasets from those observed in the extant taxa. b) Despite these differences, the performance of TED does not suffer from increasing proportions of the morphological data evolving under a non-stationary model, with slowly-evolving characters invariably retaining their full inference power. c) Example trees demonstrating the difference between the signal in the molecular (no modification) versus the morphological partition, with the labels of the extant taxa randomized in the latter (fossil taxa not shown). The two trees were inferred from one of the simulated datasets using neighbour-joining on uncorrected p-distance matrices and plotting the taxon identities among the two trees using co-phylogenetic methods that maximise visual correspondence. d) Decreasing performance of TED under increasing portions of taxa being randomized, either changing only the labels of the fossils, only those of the extant taxa, or both. Irregularities of the histograms in cases of low performance are due to the small number of replicates and should not be overinterpreted.

A more deleterious situation arose when the morphology partition reflected an underlying tree that was increasingly different of the species phylogeny (Figs 5c and d). Such a situation might result for instance in cases of extensive convergence due to ecological similarities that lead to an extensive mismatch between the trees implied by the morphological versus the molecular data partitions (Fig. 5c). If such mismatch is strong, TED is bound to fail (Fig. 5d). The lower effect when fossil labels were exchanged randomly versus those of extant taxa can probably be attributed to their lower number of fossils versus extant taxa in our simulations: in the simulation trees with 12 extant taxa, only four fossils were added.

### Deviations from the morphological clock

To mimic a low temporal signal in the morphological data, we simulated trees with increasing rate variance among the branches of the trees (Fig. 6), choosing rate multipliers based on a lognormal distribution (Fig. 6a). Examples of the simulation trees resulting from these branch-length modifications are shown in Figure 6b and range from clock-like to strongly non-clock-like (beware that sampled fossils are also shown in these trees and were also subjected to branch length transformations). When running separate relaxed-clock models with independent gamma-rates for the morphology versus molecular partitions, the analyses assuming a separate clock for the two data types correctly identified higher clock variance for the former than the latter (Fig. 6c). Nevertheless, the analyses under the wrong clock models, i.e., the strict clock or a single relaxed clock for the combined morphological and molecular partitions, did not perform significantly worse than the two-clock approach. Indeed, our analyses indicated that the effect of increasing variance in the morphological clock only has a negative impact on TED when it is extreme (at a variance of 10 in our simulations), and even then only in small trees. Under extreme variation in the clock rate among branches, even the strict-clock model still performed very well for trees with 48 extant and 16 fossil taxa, despite the temporal signal not being obvious anymore at all when looking for the fossil branches in the underlying simulation trees (Fig. 6b).

**Figure 6.**
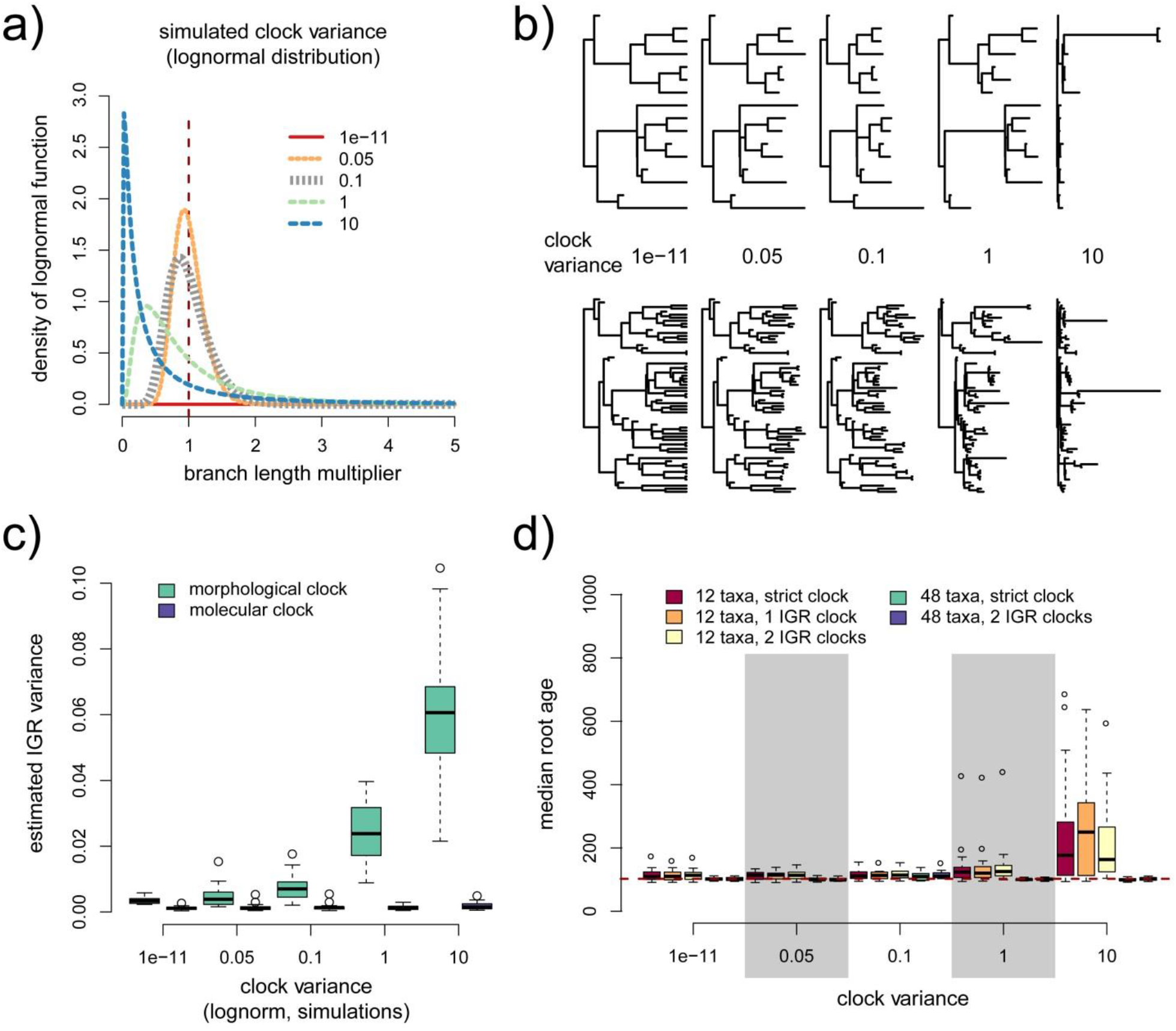
Decreasing the clock-likeness of the morphological data partition. a) Lognormal distributions with mean = 1.0 and varying variance, from which branch length multipliers were drawn. b) Examples of resulting trees with increasing clock variance, both for datasets with 12 extant and four fossil and 48 extant and 16 fossil taxa. c) Clock variance as inferred by MrBayes from these datasets under the independent gamma rates (IGR) model with two separate clocks for the morphological versus molecular data partitions, respectively. The clock variance was correctly identified as being close to zero for the 1,000-character molecular partition. d) Performance of TED when increasing the variance in the morphological clock, under a strict clock, single relaxed clock, and two relaxed clock models, for two different tree sizes. Irregularities of the histograms in cases of low performance are due to the small number of replicates and should not be overinterpreted.

### Fossil character sampling

To mimic the incomplete preservation of fossils, we removed the information in an increasing number of characters for the fossils, but not the extant taxa (Fig. 7). The number of characters sampled for the fossils was varied from five to 90, with missing characters either randomly chosen for each fossil taxon, or clustered in the fossils, so that in the extreme case, the fossils were all missing the same characters. For the smaller trees with 12 extant and 4 fossil taxa, between 20 and 50 characters were necessary to achieve robust results, while already five characters per fossil were sufficient in the larger trees of 48 extant and 16 fossil taxa. Biases in the way the characters were sampled in the fossils, i.e., if missing data was clustered among characters or not, had a negligible impact.

**Figure 7.**
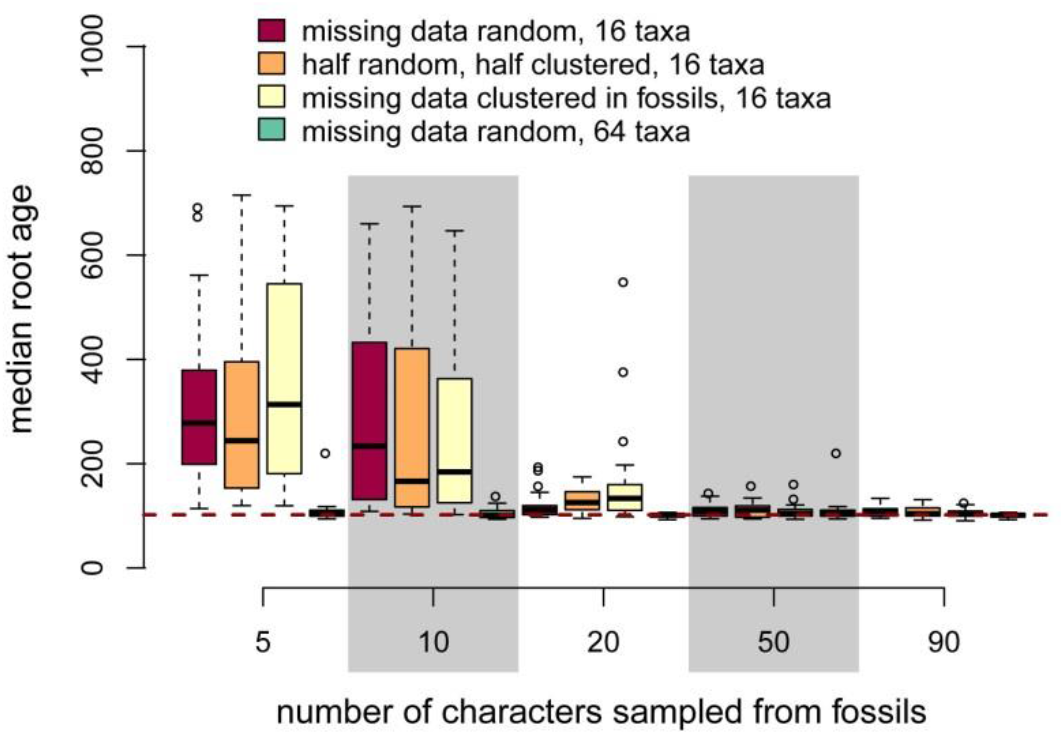
Incompleteness of character sampling in the fossils. The number of characters sampled per fossil was increased from five to 90 (out of 100 morphological characters sampled for the extant taxa), with different clustering of the missing data and two different tree sizes. Irregularities of the histograms in cases of low performance are due to the small number of replicates and should not be overinterpreted.

### Temporal and phylogenetic clustering in fossil sampling

The sampling of fossils for TED analyses is usually rather restricted by the availability of reasonably-well preserved specimens. In order to analyse the impact of different fossil sampling strategies, we assessed both a limited temporal and a limited phylogenetic scope of fossil representation in a dataset. The phylogenetic clustering of fossils had a negligible impact in our simulations (Figs 8a and b), and even very small numbers of fossils (in the extreme case just one or two) were sufficient to inform TED if they included very old fossils as well (Fig. 8b). In contrast, the inclusion of only young fossils (Figs 8c to f) turned out to be detrimental for performance, and even increasing the number of fossils to six (with 12 extant taxa in the tree) did not alleviate that effect (Fig. 8d). This was even the case for the larger trees with 48 extant and 16 fossil taxa (Fig. 8f), but to a lesser extent; when the oldest fossil was at least 40 Ma old, TED performed reasonably well in the larger trees. However, if only fossils younger than 15 Ma were sampled, not even short subtending branches and extensive taxon sampling was sufficient to make up for the lack of temporal depth. The effect of increasing the lengths of the branches leading to fossil taxa is evident at least in the smaller trees (Fig. 8f): when the fossil branches are very long (30 Myr), performance was low even if one of the oldest fossil was from 65 Ma.

**Figure 8.**
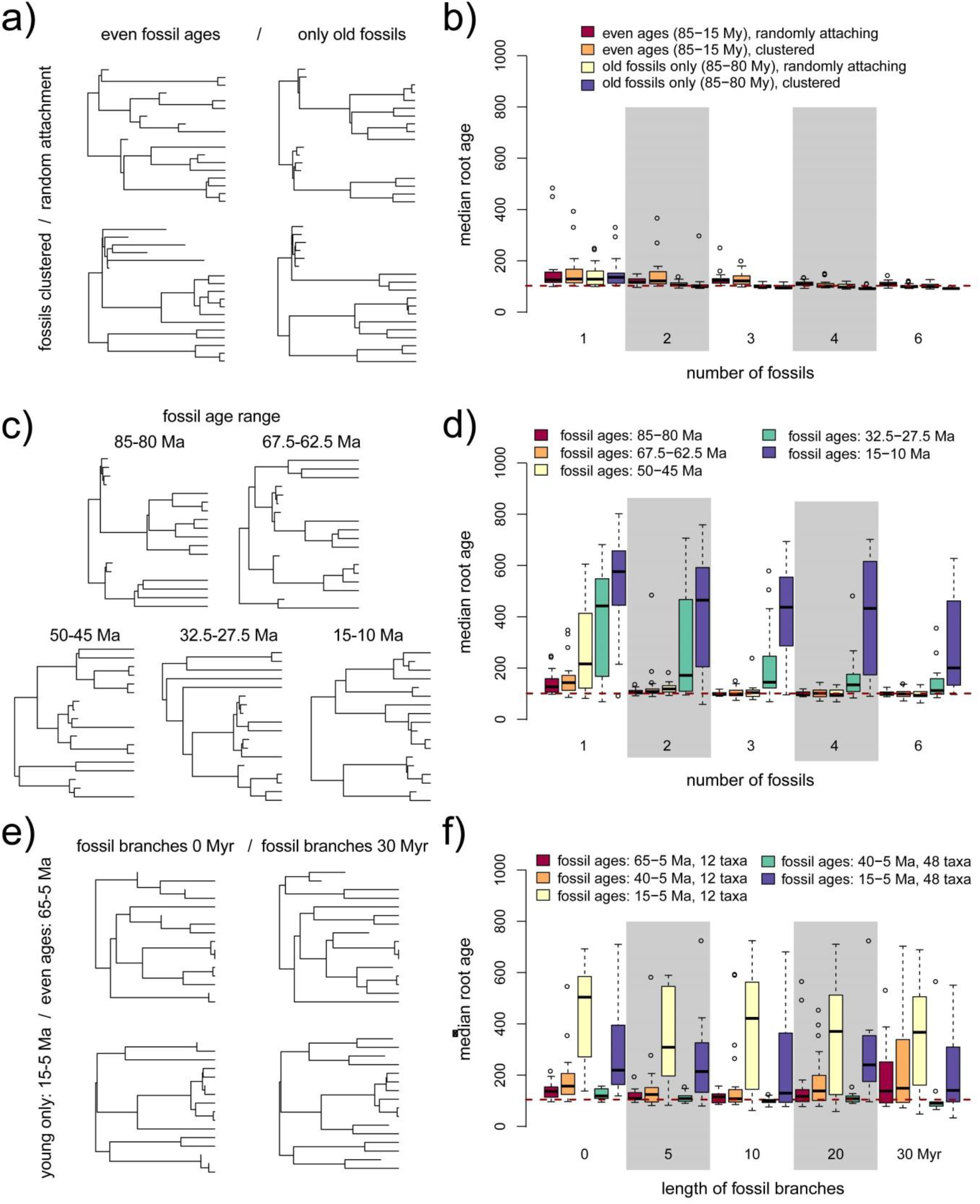
Fossil sampling strategies with respect to time and phylogenetic clustering. a) Depiction of sampling strategy: fossils were sampled either over a wide (85–15 Ma) or a narrow (85–80 Ma) age interval, keeping the oldest fossil age constant. Fossils either attached randomly to any of the branches present at the respective time (with terminal fossil branches being 5 Myr in length), or all but the first placed fossil attached to other fossil branches (phylogenetic clustering of fossils). b) Age estimates from 20 replicates with trees including 12 extant taxa and varying numbers of fossils. c) Variation of the fossil age range, with interval widths always 5 Myr and oldest fossil age decreasing from 85 Ma to 15 Ma. d) Age estimates from 20 replicates with trees including 12 extant taxa and varying numbers of fossils. e) Simulating increasing lengths of the terminal branches that lead to fossil taxa (0 to 30 Myr), for different fossil age ranges. Note that longer terminal fossil branches lead to older attachment points to the phylogeny, as the fossil ages are kept constant within replicate. f) Age estimates resulting from three different fossil age ranges and two different tree sizes (12 extant taxa + 4 fossils and 48 extant taxa + 16 fossils). Irregularities of the histograms in cases of low performance are due to the small number of replicates and should not be overinterpreted.

### Evolutionary rates in empirical datasets

Because we found that the evolutionary rates of morphological characters largely determine the impact of model mismatch on TED analyses, we estimated site rates from empirical datasets that have been compiled for the purpose of TED. Among the six empirical datasets examined (Beck and Lee 2014; Close, et al. 2016; Herrera and Davolos 2016; Kealy and Beck 2017; Lee 2016; Ronquist, et al. 2012a), we found that the vast majority of the morphological characters evolve under near-optimal evolutionary rates (0.01 to about 0.5 per root-tip distance), while none or fewer than 5% of characters have a rate which is higher than 2.0 (Fig. 9). This was true across different taxonomic groups examined, hymenopteran insects (Fig. 9a), pufferfishes (Fig. 9b), and four different mammal groups (Figs 9c-f), and regardless of how the authors of these datasets judged the credibility of the obtained age estimates. The only exception was the dataset on dasyuromorphian marsupials (Kealy and Beck 2017) which contained 8.6% of characters evolving at rates higher than 2.0 (for this dataset, the authors preferred a combined TED and node dating approach which resulted in younger age estimates).

**Figure 9.**
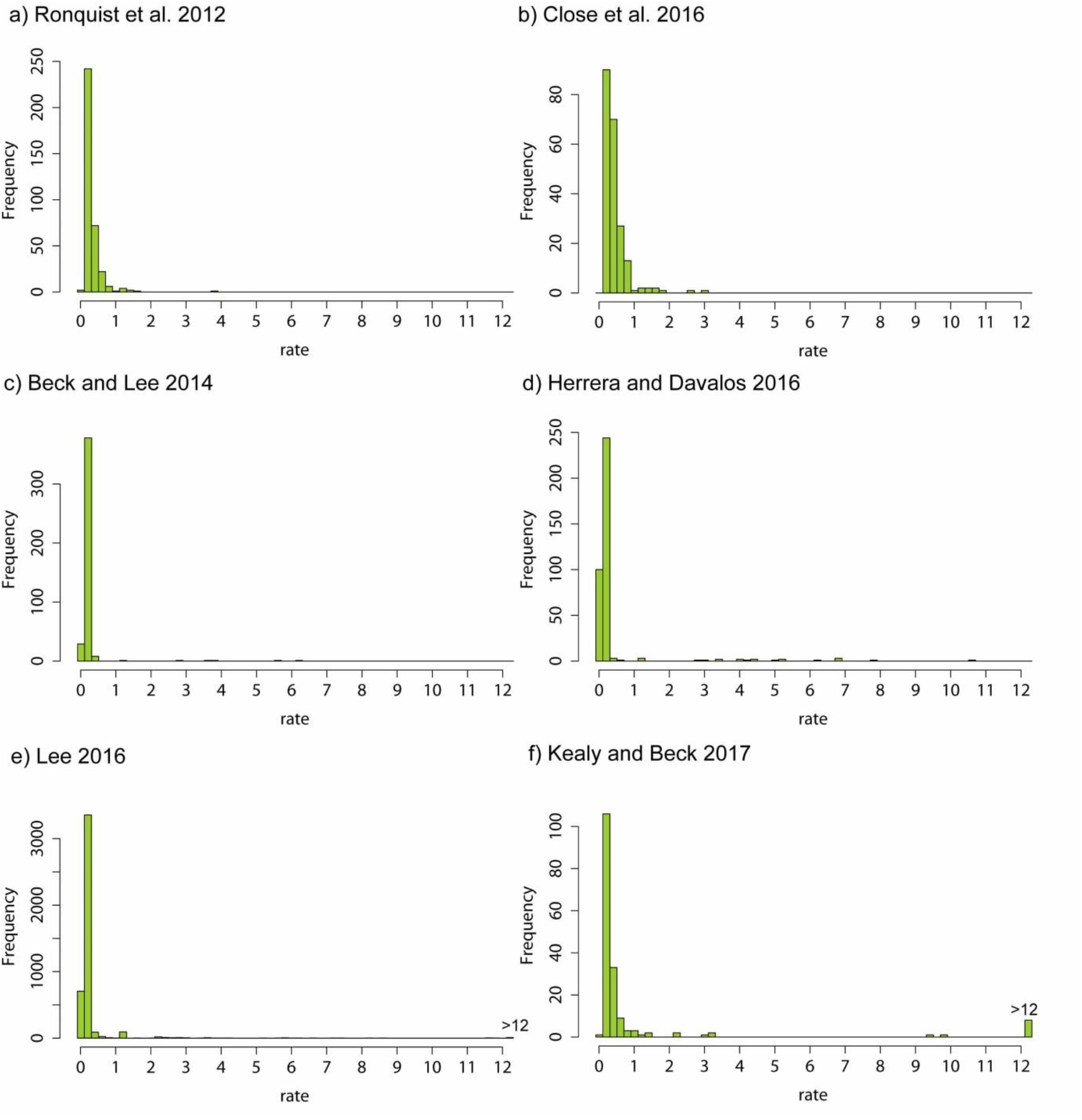
Evolutionary rates of morphological characters in six empirical datasets. Rates are given in expected changes per root-to-tip distance in the resulting tree. The data stem from six total-evidence dating studies that cover hymenopteran insects (a), pufferfishes (b), and different mammal datasets (c-f). If present, characters evolving at a rate of more than 12 changes from root to tip were summarized in a single bin (“>12”).

## Discussion

### Mismatch of the Markov model is mostly unproblematic in total-evidence dating

The large robustness of TED against misspecifications of character states and transition rates is maybe the least expected result of this study. We demonstrate that even large differences between underlying evolutionary pathways on one hand and observed character states and assumed rate models on the other have a negligible impact on the accuracy of the analysis when evolutionary rates are low. Comparisons with a few empirical datasets show that morphology partitions regularly consist of mostly very slowly evolving characters, for which we can assume that model mismatch is negligible.

A similar observation has been made already in the context of the molecular substitution model, where mismatch also had only a minor impact on divergence time estimates (but in the context of node dating; Schenk and Hufford 2010). With respect to mismatch of morphology models, only the ascertainment bias inherent in morphological dataset has been studied in the context of dating (Luo, et al. 2019; Matzke and Irmis 2018), and we here could confirm their conclusion that this bias does not have a major impact on TED performance. The low impact of misspecification of the Markov model on TED at low evolutionary rates, including misspecification of states and rates, has to our knowledge not been noted before. The reason for it might lie in the fact that for slowly-evolving characters, multiple changes along a single branch are very rare, and morphological branch lengths are informed rather by the number of characters that change along a particular branch than by an estimate of multiple changes in a single character. Misspecification of the Markov model as it results from an imperfect understanding of the molecular and developmental basis of observed morphological variation might thus be far less relevant for TED performance than previously assumed (Lee, et al. 2014). This is the case both for trees with a small number of terminals and for larger trees, despite the fact that the power to distinguish between the correct and incorrect models of evolution can be expected to increase in the latter; maybe the negative effects of the model mismatch are balanced out by the temporal information stemming from additional fossils. This result is very reassuring, given our still patchy knowledge of the relationships between genome, environment and phenotype and resulting difficulties in designing realistic models of morphological evolution.

The six empirical studies that we have chosen to estimate evolutionary rates of their morphological partitions might or might not reflect a random sample. We focussed on studies that used datasets chosen specifically for the purpose of TED (Beck and Lee 2014; Herrera and Davolos 2016; Kealy and Beck 2017; Lee 2016; Ronquist, et al. 2012a), including both presumably well-behaved datasets and some for which the authors of the study suspected a low performance of TED (based mostly on comparisons with the fossil record of a group). It remains to be shown whether morphological datasets in general contain only few fast-evolving characters; this at least seems likely, given that morphologists often mostly choose slow characters which are believed to be more informative for phylogenetic reconstruction due to low levels of homoplasy. If morphological data contain more slowly-evolving characters than molecular data (while typically being devoid of constant characters), it can be assumed that mismatch of the Markov model would be less problematic, even if said mismatch has been found to be quite pronounced in morphological datasets (Goloboff, et al. in press).

We assumed in our simulations that model mismatch affects all branches of the tree equally, and we have not investigated whether clade-specific mismatches would lower the performance of TED. Except for a set of functionally linked characters, it is difficult to think of a scenario in which a particular clade would suffer from more severe model mismatch than the others, except maybe for groups which have undergone drastic miniaturisation or paedomorphosis and thus exhibit an impoverished character state space in comparison to the remaining taxa in the tree (Bleidorn 2007; Wiens, et al. 2005). Such a bias would be reflected in shorter morphological branch lengths in that clade, which is equivalent to a slow-down of the morphological clock – an aspect we did cover in our analyses of clocklikeness (see below). Nevertheless, heterogeneous model mismatch across the tree deserves further attention.

### Beware of asymmetric state frequencies and non-phylogenetic signal in morphological data

The only aspect of mismatch of the Markov model that had a severe impact in our analyses was the misspecification of the asymmetry in state frequencies. This is a very common aspect of morphological datasets, where for instance the “absent” state of a character is regularly overrepresented (Wright, et al. 2016). While the modelling of asymmetry is straightforward in molecular data, where state labels have biological meaning (i.e., they represent nucleotides or amino acids with common properties), morphological characters employ arbitrary state labels, with the meaning of an individual state label changing between characters; Lewis (2001) already proposed a way to circumvent this issue by assuming a distribution of asymmetry across characters modelled in a similar way to ASRV (Yang 1996). We here used a hyperprior to model differing strengths of asymmetry across characters, which proved to be insufficient to overcome the bias introduced by strong asymmetry in the smaller tree with 16 taxa, while behaving very well in the case in our larger tree of 64 taxa. It can be assumed that the additional information about morphological evolution in the larger tree was necessary to model asymmetry accurately; it remains to be shown how the model behaves on trees of intermediate size. Given that we assumed very strong asymmetry in all the characters, while empirical datasets will probably include at least some nearly symmetrical characters as well, we probably covered the most severe effects that asymmetry can have on TED performance in our simulations. In any case, this aspect of morphological data certainly deserves further attention, also with respect of mismatch of the (symmetric) Dirichlet distribution used to model across-character asymmetry in state frequencies.

Another aspect of morphological data that we identified as detrimental for TED is rampant non-phylogenetic signal, as it could result from functional or structural convergence or parallelism (i.e., morphology reflecting ecology or life history rather than phylogenetic relationships; Ronquist, et al. 2016). The impact of non-phylogenetic signal was most severe when both fossil and extant taxa were simulated to evolve on the wrong tree for the morphological partition. While we investigated different portions of taxa being randomized, reflecting a scenario in which varying numbers of unrelated terminals are grouped together for the entire morphological partition, most empirical datasets will consist of a mixture of characters with either phylogenetic or ecological signal. Our simulations can thus be viewed as rather extreme in that the entire morphological partition was concerned; it remains to be shown how much phylogenetic versus conflicting signal is needed to mislead TED. In any case, strong conflict between the topology supported by the molecular versus the morphological partition of a dataset should be taken as indication that age estimates based on TED might be misleading (Ronquist, et al. 2016).

### Morphological clock and clock-model mismatch

Our simulations demonstrate that even a surprisingly weak morphological clock can provide sufficient time information for TED, especially in trees with a few dozen or more extant taxa and a good number of fossils. Clock-likeness is often lower in morphological than in molecular datasets (Beck and Lee 2014; Ronquist, et al. 2012a), but the variance among branch rates observed in these morphological datasets was in all these cases well within the range covered by our simulations (see also Fig. 6b). Individual branches that were subject to a strong speed-up in morphological evolution do not seem to have a strongly negative impact, probably because there are still enough fossils on other branches that evolve at a rate closer to the overall morpho-clock rate to inform the correct timing of events.

A similarly low impact of among-branch rate variation on TED was already found in a previous simulation study which investigated the method under the FBD tree prior (Luo, et al. 2019). Our simulations have used more extreme values for the variation of the morphological clock than this previous study, and still we found that ALRV is not a major issue for TED analyses. What might surprise even more, not only in the context of morphological but also of molecular clock analyses, is the observation that mismatch of the clock model did not have any impact on the resulting age estimates, with the strict-clock model performing as well as the (incorrect) single-clock model and the (correct) two-clock model, which estimated relaxed-clock branch lengths separately for the molecular and the morphological partition (Fig. 6d). This is certainly worth investigating further, especially since relaxed-clock models regularly cause problems with convergence in Bayesian MCMC analyses, as we also observed here: we had to increase the run-time by a factor two to three in order to obtain sufficient convergence on the clock-rate parameter under relaxed in comparison to strict-clock analyses. We also observed that convergence was crucial, with highly misleading age-estimates resulting from MCMC runs that were aborted to early. MCMC convergence is far easier to achieve under the much simpler strict-clock model, while our simulations suggest that no power is lost for estimating divergence times, at least under the type of clock variance simulated here; additional simulations studies are needed to show whether this assertion also holds for other types of clock variance among lineages.

A related characteristic of morphological data has received increased attention recently: heterotachy or the lack of a shared set of branch lengths across characters (“no common mechanism”; Goloboff, et al. 2018; Goloboff, et al. in press; O’Reilly, et al. 2016; O’Reilly, et al. 2018). In our simulations, we approached heterotachy by randomly drawing different rate multipliers for each branch, in the most extreme case using a separate set of multipliers for every single character; even though this resulted in large differences among underlying branch length distributions across characters, we still assumed some connection of the time lengths of a branch and its evolutionary length for morphological characters. It is thus maybe not surprising that heterotachy improved TED performance in our case: the different rate multipliers acting on each character effectively cancelled each other out, leading to very good performance of TED even in cases of high variance in rate multipliers or per-character clocks. Whether these simulations are relevant in reality depends on whether there is any correlation at all between morphological branch lengths and time, or, by proxy, between morphological and molecular branch lengths. Current consensus in the literature (Davies and Savolainen 2006; Halliday, et al. 2019; Omland 1997; Ronquist, et al. 2012a) ascertains that this is mostly the case (but see Bromham, et al. 2002). But of course datasets might differ in that respect, and TED studies should test their morphological partition for clocklikeness before deciding to rely solely on morphological branch lengths for divergence time estimation.

### Extensive, empirical study of the fossil record is crucial

While mismatch of the morphology model appears to be a minor issue under many circumstances, our simulations underline the central role of an extensive sampling of fossils in TED studies, especially of older ones. A similar result was found by Luo et al. (2019) and O’Reilly and Donoghue (2019), who already emphasized the vital role of extensive fossil sampling. In contrast to node dating, which is based largely on the oldest representative of each clade, the entire fossil record can be used for TED (Ronquist, et al. 2012a). Even highly incomplete fossils can and should be included, no matter if they can be placed in the phylogeny with any certainty. The high instability of placement especially of fragmentary fossils might deter researchers from coding them for morphology, but previous TED analyses have demonstrated that even fossils coded for very few morphological characters potentially contribute to the precision of divergence time estimates. In our simulations, as few as five characters coded per fossil were sufficient to inform robust age estimates in the case of the larger tree with 48 extant and 16 fossil taxa. This suggests that a lower number of characters per fossil could potentially be compensated for by sampling more fossils, with the placement certainty of the individual fossil playing only a minor role for TED performance. Increased fossil sampling could also partly offset a low temporal scope in our simulations, except if age ranges only covered less than a third of the depth of the tree. TED studies that recover root ages that are several times higher than the oldest included fossils should thus be met with caution, as the temporal information has likely not been sufficient. When designing TED studies, priority should be given to the detailed study of the available fossil information of a group.

### A plea for TED under the uniform tree prior

Our simulation study suggests that total-evidence dating is much more robust to model mismatch and low clocklikeness of morphological data than previously thought, and that it can work even with highly incomplete fossils. It establishes TED under the uniform tree prior as a valid alternative to the more commonly used combined approaches of TED with the FBD tree prior and/or node age priors, especially in cases where the fossil record is either scarce or irregular. It remains to be shown whether previous failures of TED on particular datasets were due to biases in the morphological data that were more extreme than those mimicked in our simulations, or if the type of bias was not covered here at all, or if other aspects of the analyses were inadequate, such as fossil sampling or node-age or clock-rate priors. In any case, where strong enough, the morphological clock could act as an ideal complement to the molecular clock that allows inclusion of fossil taxa. It would allow us to avoid relying on uncertain secondary interpretations of the fossil record, like they are used in node dating, or on assumptions about the fossilization process, which might bias fossilized-birth-death analyses. It remains to be demonstrated what proportion of morphological datasets meet these requirements, and whether the morphological clock can be strengthened by improved character concepts and morphology models. However, our results clearly show that the effort involved in coding morphological characters and implementing fossils in TED analyses can be repaid with a better understanding of the timeline of life.

## Material and methods

Simulations were performed in R (R Core Team 2014), making use of the packages “phangorn” (Schliep 2011) and “TreeSim” (Stadler 2011). Scripts are available as Supplementary Material S1. A number of shell scripts were used to set up an analysis pipeline, including the generation of input files, parallel submission of Bayesian analyses and summarizing of the results. Phylogenetic analyses were performed in MrBayes 3.2 (Ronquist, et al. 2012b) on UBELIX (http://www.id.unibe.ch/hpc), the HPC cluster at the University of Bern, Switzerland.

### Simulation trees

We simulated phylogenetic trees with 12 and sometimes also with 48 extant taxa (Fig. 2a), respectively, under a pure birth process, with the birth rate optimized for the number of tips (birth rate = ln (number of taxa / 2) / tree age). These trees were chosen deliberately to be rather easy to resolve, instead of using a combination of birth and death rates that would lead to short basal branches in the tree (tree imbalance as a factor influencing TED performance has been covered elsewhere: Duchêne, et al. 2015). The trees were then stretched so that their root age was equal to 100 My. For each simulation condition, 20 replicate trees were simulated in the case of 12 taxa and 10 in the case of 48 taxa.

Instead of applying a model of fossilization and fossil recovery to simulate fossil placements (Luo, et al. 2019; O’Reilly and Donoghue 2019; Stadler 2010), we decided to use a pattern-based approach that would provide us with full control over fossil sampling conditions. In most analyses, we followed the following approach (but see below for changes to this standard setting): we chose sampled fossils evenly through time, with the oldest being 85 My and the youngest 15 My old. In the case of 12 extant taxa, we added four fossils, while the trees of 48 extant taxa contained 16 fossils (1/3 of the extant taxa). The terminal branches leading to the fossils were 5 My long and were attached randomly to one of the branches present at the respective time, starting with the oldest fossil. In the larger tree of 48 taxa, it was thus possible that a fossil would attach to a branch leading to a fossil and not to any extant taxa. Examples of simulation trees are shown in Figure 2a; they reflect a presumably favourable tree shape and fossil sampling, with both a good temporal and phylogenetic spread and rather short terminal branches leading to the fossil taxa.

### Simulation of molecular characters

For all extant taxa, we simulated nucleotide sequences of length 1,000 bp under the Jukes-Cantor model, a strict clock, and with a clock rate of 0.0025 substitutions per Myr for all sites. This rate translates into 0.25 expected substitutions between root and tip on our tree, which is 100 Myr deep, a rate that has been found optimal for topology inference in a recent simulation study (Klopfstein, et al. 2017). For the nucleotide data, we always used the same model for simulation as for analysis, ensuring straightforward recovery of the relationships among the extant taxa; the molecular data thus in fact almost acts as a full constraint on the topology and molecular branch lengths (see results section).

For all analyses, we set the tree age prior to a uniform distribution between zero and 1,000 Ma, and this prior was recovered very well in the eight independent test runs without data (Fig. 9a). However, in dating analyses, another prior might influence the effective prior on node ages: the prior on the clock rate. We aimed for a rather flat distribution, but one that invokes a clear bias away from the true value of 100 Ma. We thus chose to set the clockrate prior to a log-normal distribution with a mean that matches the true value (μ = −11.98293, σ = 3.461638, mean = 0.0025). Its variance was chosen in a way that, in the absence of temporal information (i.e., when running without fossils but with the extant taxa), creates an expected root age estimate under our simulation settings that is around 575 Ma (Fig. 9b), with the posterior probability of it being below 120 Ma below 5% (both for trees with 12 and with 48 extant taxa). We thus know what to expect when total-evidence dating fails: root age estimates will tend towards too high values.

**Figure 9.**
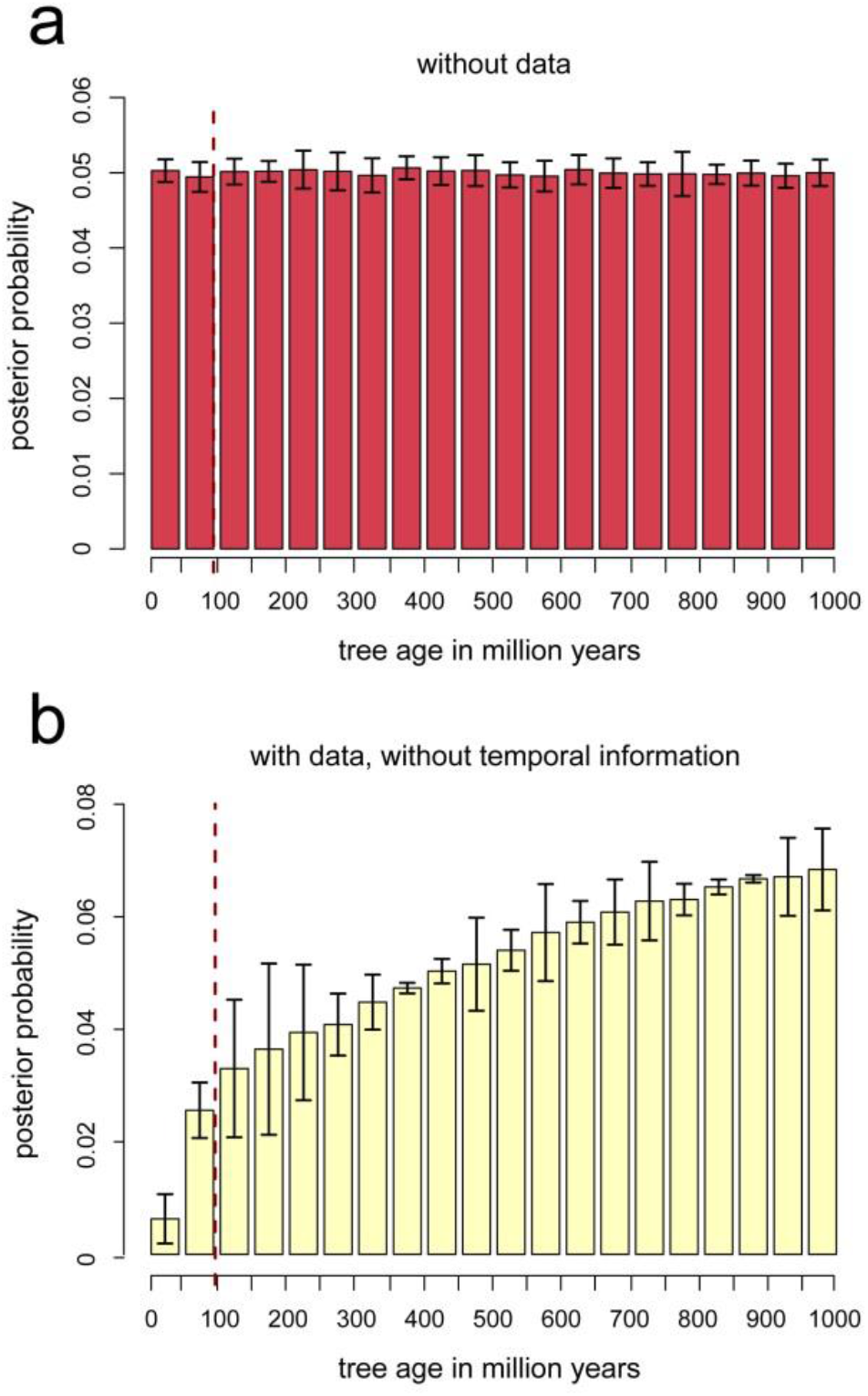
Effect of tree age and clock rate priors on posterior age estimates in trees with 12 extant taxa. a) Results from eight independent runs without data, recovering the tree age prior, a uniform distribution between zero and 1,000 Ma. b) Posterior distribution implied by both the tree age prior and by the prior on the clock rate, as retrieved in two independent runs with molecular and morphological data, but without the fossils, thus removing all time information. The log-normal clock rate prior was chosen so that it implies a much higher tree age than the true one used in the simulations (100 Ma, indicated by the vertical dashed line). Posterior probabilities of the tree age being recovered as below 120 Ma were below 5%.

### Morphology model and model mismatch

To mimic the morphology partition, we simulated 100 binary, stationary characters with equal state frequencies, in most cases sampling only variable characters, as is the rule for morphological datasets (corresponding to the Mkv model with k = 2; Lewis 2001). Fifty of these 100 characters were sampled for the fossils, chosen randomly and independently for each taxon in most analyses (but see below). Except for the analysis in which we addressed the impact of the evolutionary rate of the morphological characters on TED performance, we used a highly informative rate of 0.0025 changes per Myr for morphology, which is identical to the one used for the molecular characters. We then varied different aspects of this basic model for the simulations, but not for the analyses, creating different flavours and extents of model mismatch for the morphological partition. First, we addressed ascertainment bias by sampling all characters, only variable characters, or only parsimony-informative characters. At the same time, we varied the evolutionary rate of the morphology partition between 0.05 and 10 substitutions between root and tips (i.e., 0.0005 to 0.1 changes per My), with equal rates across characters.

The tempo of evolution can vary strongly between morphological characters; we thus simulated increasing extents of among-character rate variation (ACRV). While keeping the average rate across characters at 0.0025 changes per Myr, we first used a discretised gamma distribution with four rate categories for simulation, varying the shape parameter between 100,000 (almost equal rates among characters) and 0.01 (very strong ACRV). This data was then analysed under the gamma model with the distribution’s parameter estimated from the data (no model mismatch), and under a model that assumes equal rates (creating model mismatch). To assess whether the flavour of ACRV model mismatch had an impact, we also simulated data assuming a distribution with two peaks: half of the characters were simulated under the lowest of the four rate categories and the other half under the highest one. This “bimodal” data was analysed under the discretised gamma distribution.

One of the main concerns in morphological phylogenetics, both for character coding and analysis model, is the mismatch between the observed character states and the underlying evolutionary process. To mimic a case where the states have been identified correctly, but transitions between them were misspecified, we simulated data under an “ordered” model with three states, in which only transitions between adjacent states are allowed. For example, if the states are denoted as 0, 1 and 2, the transition between 0 and 2 could not go directly, but would need to pass through state 1. In the analyses, we then assumed that all transitions are equally likely (standard Mkv model). We varied the number of ordered states (and thus the extent of model mismatch) between 2 and 10 states.

To reflect a type of model mismatch where the states have been misidentified, we simulated data under a 10-state ordered model, with characters transitioning only between adjacent states. We then randomly chose for each state (from state labels 0 to 9) whether to transform it into the observed state 0 or 1, effectively creating a binary character. The transformation was also done for less severe losses of information, transforming randomly into 3, 6, or 10 states, respectively, always using an unordered model for analysis. This set-up was chosen to reflect a situation in which multiple, unrelated underlying genetic and/or developmental mechanisms result in the same observable morphological character state. To create an additional type of information loss, we also varied the number of underlying states using the same values as above and always transforming them into a two-state character.

The standard Mk model assumes equal state frequencies (and thus transition rates; Lewis 2001), which is not matched by many morphological datasets. We simulated such asymmetry by decreasing the frequency of state 1 in binary characters from 0.5 (no asymmetry) to 0.1 (strong asymmetry) under different rates of morphological evolution. Analyses were performed under the symmetric model.

Many morphological character systems follow a directional pattern across parts of the tree of life, be it due to positive selection or biased starting conditions (Gould 1996; Klopfstein, et al. 2015). To mimic increasing levels of directionality, we varied the proportion of characters in the dataset that followed a non-stationary pattern between zero and one by adjusting the state frequencies at the root. The non-stationary characters would then all start out at state 0, while stationary characters would show an equal probability of the two states at the root of the tree.

### Morphological clock mismatch

To study the effect of a lower clock-likeness of the morphological data partition, we modified our simulation trees to mimic increasing amounts of deviation from the strict clock. Branch length multipliers were drawn randomly for each branch (including those leading to fossils) from a lognormal distribution with a mean of one and increasing variance from zero to ten. This produced trees with zero among-lineage rate variation (ALRV) to trees with extensive ALRV (Fig. 6b), on which the morphological data was then simulated. The molecular partition was assumed to evolve in a strictly clock-like fashion. For the analyses, we then compared approaches assuming a strict clock for both partitions to a global relaxed clock and to a separate relaxed clock for the morphology and the molecular partition.

### Sampling of fossils and fossil characters

Morphological characters in fossils are more difficult to sample than in extant taxa due to incomplete preservation. We thus varied the number of fossil characters sampled from five to 100, either subsampling them randomly for each fossil, keeping the same characters present or absent in all fossils, or a mix of these two sampling strategies.

To assess the impact of fossil sampling, we first used a wide (85–15 Ma) and a narrow (85 – 80 Ma) temporal interval, in each case either sampling fossils that were attaching randomly to the branches of the tree present at the time, or in a clustered manner, so that all but one fossil branches attached to each other (“phylo-temporal clustering”, see Tong, et al. 2018). The narrow interval of 5 Myr was then shifted over time, from 85–80 to 15–10 Ma, to assess the temporal depth needed in total-evidence dating analyses. To study the effect of the length of the terminal branches leading to the fossils and the interaction of this length with the temporal scope in fossil sampling, we increased terminal branches from length zero (mimicking sampling true ancestors of extant taxa) to 30 Myr, for fossil ages distributed evenly between 5 and 15, 40, or 65 Ma, respectively.

### Estimating evolutionary rates in empirical datasets

In order to validate our simulation results, we estimated evolutionary rates of morphological characters from six empirical dataset that have been compiled for total-evidence dating. The chosen datasets cover insects (Ronquist, et al. 2012a), fishes (Close, et al. 2016), and four different mammal groups (Beck and Lee 2014; Herrera and Davolos 2016; Kealy and Beck 2017; Lee 2016). We estimated evolutionary rates of each character individually in R (R Core Team 2014) using the function “ACE” from the “ape” package (Paradis, et al. 2004) under the equal rates model. This function takes a fully resolved rooted tree and a character matrix as input. For all datasets except for Beck and Lee (2014), we used one of the publicly available trees: a maximum clade credibility tree (Kealy and Beck 2017), a maximum sampled-ancestor clade credibility tree (Close, et al. 2016), a maximum likelihood tree (Herrera and Davolos 2016), or the tree with the highest likelihood from a set of Bayesian posterior trees (Lee 2016; Ronquist, et al. 2012a). An appropriate tree was not available for the Beck and Lee (2014) study; we thus re-run their Bayesian analysis with the same settings and selected the tree with the highest likelihood. Before site-rate estimation, we excluded the tips with missing data for the character in question.

## Acknowledgments

This study was funded by the Swiss National Science Foundation (grant PZ00P3_154791 to SK).

